# A maximum mean discrepancy approach reveals subtle changes in *α*-synuclein dynamics

**DOI:** 10.1101/2022.04.11.487825

**Authors:** Hippolyte Verdier, François Laurent, Alhassan Cassé, Christian L. Vestergaard, Christian G. Specht, Jean-Baptiste Masson

## Abstract

Numerous models have been developed to account for the complex properties of the random walks of biomolecules. However, when analysing experimental data, conditions are rarely met to ensure model identification. The dynamics may simultaneously be influenced by spatial and temporal heterogeneities of the environment, out-of-equilibrium fluxes and conformal changes of the tracked molecules. Recorded trajectories are often too short to reliably discern such multi-scale dynamics, which precludes unambiguous assessment of the type of random walk and its parameters. Furthermore, the motion of biomolecules may not be well described by a single, canonical random walk model. Here, we develop a methodology for comparing biomolecule dynamics observed in different experimental conditions without beforehand identifying the model generating the recorded random walks. We introduce a two-step statistical testing scheme. We first use simulation-based inference to train a graph neural network to learn a fixed-length latent representation of recorded random walks. As a second step, we use a maximum mean discrepancy statistical test on the vectors of learnt features to compare biological conditions. This procedure allows us to characterise sets of random walks regardless of their generating models. We initially tested our approach on numerical trajectories. We then demonstrated its ability to detect changes in *α*-synuclein dynamics at synapses in cultured cortical neurons in response to membrane depolarisation. Using our methodology, we identify the domains in the latent space where the variations between conditions are the most significant, which provides a way of interpreting the detected differences in terms of single trajectory characteristics. Our data show that changes in *α*-synuclein dynamics between the chosen conditions are largely driven by increased protein mobility in the depolarised state.

**Author summary:** The continuous refinement of methods for single molecule tracking in live cells advance our understanding of how biomolecules move inside cells. Analysing the trajectories of single molecules is complicated by their highly erratic and noisy nature and thus requires the use of statistical models of their motion. However, it is often not possible to unambiguously determine a model from a set of short and noisy trajectories. Furthermore, the heterogeneous nature of the cellular environment means that the molecules’ motion is often not properly described by a single model. In this paper we develop a new statistical testing scheme to detect changes in biomolecule dynamics within organelles without needing to identify a model of their motion. We train a graph neural network on large-scale simulations of random walks to learn a latent representation that captures relevant physical properties of a trajectory. We use a kernel-based statistical test within that latent space to compare the properties of two sets of trajectories recorded under different biological conditions. We apply our approach to detect differences in the dynamics of *α*-synuclein, a presynaptic protein, in axons and boutons during synaptic stimulation. This represents an important step towards automated single-molecule-based read-out of pharmacological action.

## Introduction

Numerical models make it possible to generate synthetic observations of biological systems across a broad range of parameters. However, the computational cost of directly using these simulations to perform statistical inference is often prohibitive [1]. The reason is that both the likelihood and evidence are intractable in most systems, and these inferences must therefore be addressed as likelihood-free simulation-based inferences [2]. The primary approach to simulation-based inference is approximate Bayesian computation (ABC) [2]. ABC relies on comparing user-defined summary statistics from experimentally recorded and simulated data using a chosen distance metric. The ever-growing amount of available data, along with recent advances in deep learning [3] make it possible to capture more and more detailed properties of experimental systems, and have thus boosted the development of simulation-based inference. We refer the interested reader to [1] for a detailed taxonomy of simulation-based inference methods.

Typical inference schemes developed to analyse biomolecule trajectories focus on estimating physical parameters such as the diffusion coefficient, the anomalous diffusion exponent, the type of random walk model, or other ad-hoc quantities measuring particular aspects of the dynamics. Here, instead of describing trajectories using a set of explicitly defined features, we rely on an *encoder* neural network, in order to characterise each trajectory by a *latent vector* of features. The goal of this encoder is to automatically learn optimised features that describe random walks beyond predefined canonical models and features. We employ the encoder network to project recorded trajectories into the fixed-dimensional latent space. We develop a statistical test on this latent space to test for differences in dynamics between two sets of trajectories. Our methodology can in particular be used to compare dynamics observed in different biological conditions and different cell organelles, by comparing the sets of latent vectors computed from trajectories observed in the respective microscopy recordings or regions of the cell. The central advantage of the testing methodology we propose is that it is not dependent on the specification and selection of a model of the recorded random walks. This enables statistically robust testing of differing biological conditions, which are likely to induce different levels of cellular heterogeneity and do not necessarily generate canonical random walks.

We train the encoder network using a simulation-based inference framework (detailed below), allowing it to provide a representation of trajectories without assuming that they are realisations of a canonical random walk model. The subsequent statistical test seeks to differentiate the distributions of latent vectors coming from different conditions or organelles (or both). It is based on the maximum mean discrepancy (MMD) test [4], which uses a kernel approach to compare two distributions. This test allows us to compare sets composed of different numbers of trajectories, and provides a means for interpreting the differences between biological conditions. Finally, we show the robustness of the approach to the intrinsic variability of biological observations, and demonstrate that the statistical differences do not stem from a single experiment, nor from an outlier composed of a minority of trajectories.

We demonstrate our methodology by studying the dynamics of *α*-synuclein inside and outside of synapses. *α*-synuclein is a small, soluble, and highly mobile protein (140 amino acid residues) that is strongly accumulated in presynaptic boutons (reviewed in [5]). Experiments based on fluorescence recovery after photobleaching [6] have shown the existence of at least two main modes of diffusion, one in which *α*-synuclein is transiently bound to synaptic vesicles in the synaptic bouton, and another in which the protein diffuses freely both in axons and in synaptic regions. The existence of an immobile population of *α*-synuclein molecules, taking the form of protein aggregates at synapses, has also been proposed [6]. In response to strong depolarising signals the bound population of *α*-synuclein dissociates from its synaptic binding sites and disperses in the neighbouring axon [7]. In agreement with these earlier studies, we found that *α*-synuclein dynamics differ between synapses and axons. Furthermore, depolarisation of the neurons shifts the relative frequency of the proteins from a less mobile to a highly mobile state, but it does not appear to induce qualitative changes in the type of diffusion dynamics the molecules follow. It is not clear what role this dynamic shift of *α*-synuclein plays in vesicle cycling and in the regulation of synaptic transmission. Single molecule based imaging in living neurons can help to address this question and yield new information about the physiological function of *α*-synuclein at synapses, as well as its involvement in pathological processes.

## Materials and methods

### Recording *α*Syn:Eos4 dynamics

#### Neuron cultures and *α*Syn:Eos4 expression

Primary murine cortical neuron cultures were prepared at embryonic day E17 as described previously [8]. Cortices were dissected, the tissue was dissociated and the cells were seeded at a concentration of 5 × 10^4^ cm^−2^ on glass coverslips that had been coated with poly-D,L-ornithine. Neurons were kept at 37^*o*^C and 5% CO_2_ in neurobasal medium supplemented with Glutamax, antibiotics and B27 (all from Gibco, Thermo Fisher Scientific), infected at day *in vitro* (DIV) 11-24 with lentivirus driving the expression of *α*-synuclein tagged at its C-terminus with the photoconvertible fluorescent protein mEos4b (*α*Syn:Eos4) under the control of a ubiquitin promotor, and used for experiments 7 days later. All cell culture and imaging experiments were conducted at the Laboratory for cellular synapse biology at IBENS (Paris). Procedures involving animals were performed according to the guidelines set out by the local veterinary and administrative authorities.

#### Single molecule localisation microscopy (SMLM)

Living neurons expressing *α*Syn:Eos4 were imaged in modified Tyrode’s solution (in [mM]: 120 NaCl, 2.5 KCl, 2 CaCl_2_, 2 MgCl_2_, 25 glucose, 5 pyruvate, 25 HEPES, adjusted to pH 7.4) at room temperature, using an inverted Nikon Eclipse Ti microscope equipped with a 100x oil objective (NA 1.49), an Andor iXon EMCCD camera (16 bit, image pixel size 160 nm), and Nikon NIS acquisition software. First, an image of the chosen field of view (average of 10 image frames taken with 100 ms exposure time) was taken in the green channel (non-converted mEos4b fluorescence) using a mercury lamp and specific excitation (485/20 nm) and emission filters (525/30 nm). This was followed by a streamed acquisition of 25 000 movie frames recorded with 15 ms exposure and Δ*t* = 15.4 ms time lapse (total duration: 6 min 25 s) in the red channel using a 561 nm laser at a nominal power of 150 mW for excitation (inclined illumination), together with pulsed activation lasers applied during the off time of the camera (405 nm, approx. 1-5 mW; 488 nm, 10 mW; 0.45 ms pulse). The red emission of the photo-converted mEos4b fluorophores was detected with a 607/36 nm filter (Fig. 1A).

**Fig 1.**
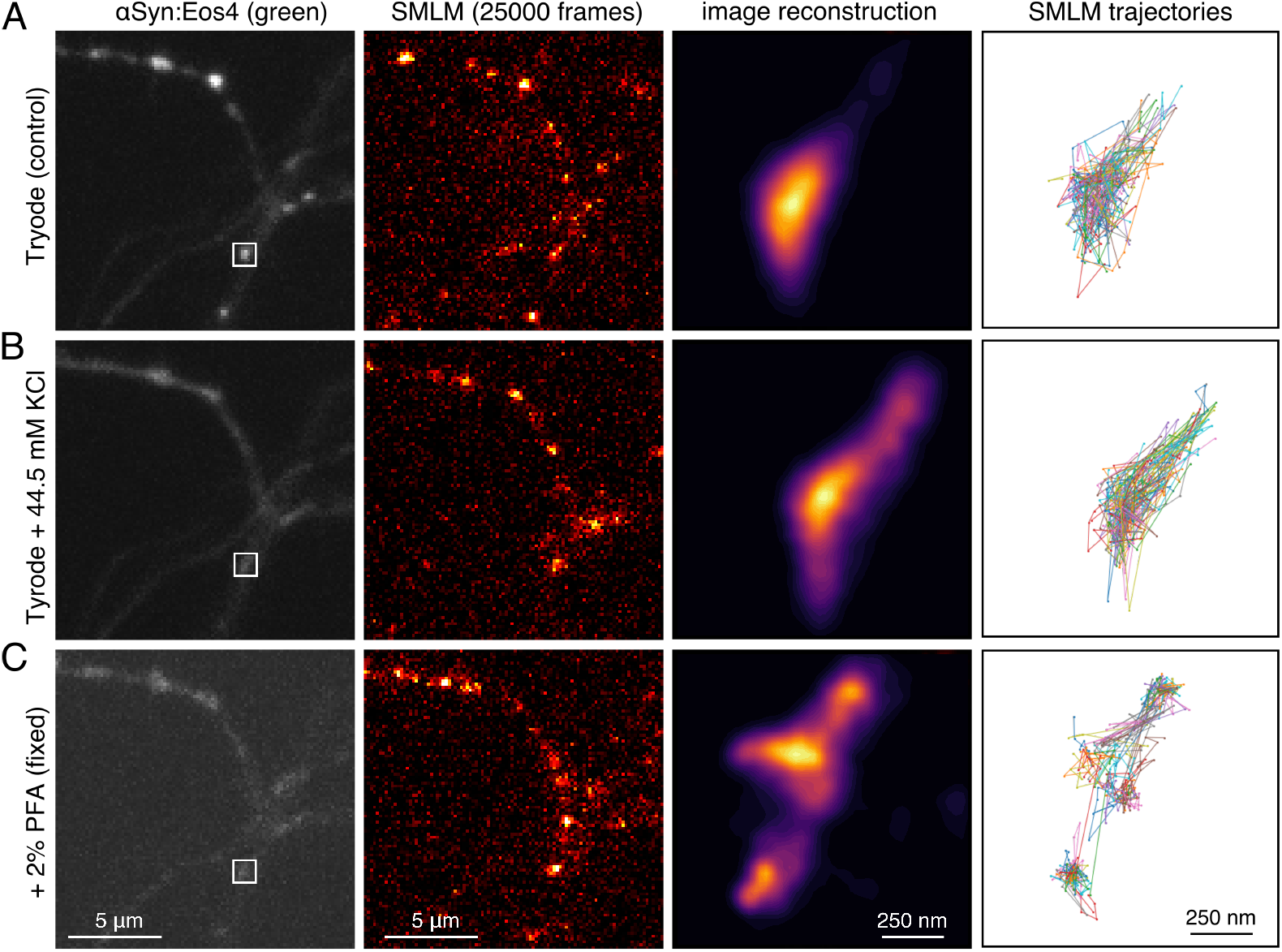
Single molecule localisation microscopy (SMLM) of *α*-synuclein in cortical neurons. (**A**) Neurons expressing *α*Syn:Eos4 were first imaged in control condition. A reference image (left panel) was taken in the green channel, followed by a SMLM movie of 25000 frames in the red channel (panel two). (**B**) A second recording of the same field of view (image and movie) were then acquired in the presence of elevated KCl concentration. Note the dispersal of *α*Syn:Eos4 in response to depolarization compared to the control (left panels). (**C**) A third image and movie were acquired after addition of 2% paraformaldehyde. The third column of images shows zoomed SMLM reconstructions of the synaptic bouton indicated in the first image. The fourth column depicts a subset of trajectories from the same synaptic terminal.

After recording of the baseline dynamics of *α*Syn:Eos4, the buffer composition was changed with the addition of Tyrode’s solution containing elevated KCl at the expense of NaCl (final concentrations in [mM]: 78 NaCl, 44.5 KCl, 2 CaCl_2_, 2 MgCl_2_, 25 glucose, 5 pyruvate, 25 HEPES, pH 7.4). This treatment causes the depolarisation of the neurons leading to the dissociation of *α*-synuclein from its binding sites in the synaptic bouton [7]. A reference image was taken in the green channel, followed by a second SMLM recording (Fig. 1B), starting approximately 7.5 min after the first acquisition. Finally, the neurons were fixed with the addition of phosphate buffer at pH 7.4 containing 4% paraformaldehyde and 1% sucrose (final concentration 2% PFA), and a third reference image (green) and SMLM movie (red channel) were acquired in the presence of the fixative (Fig. 1C).

#### Image processing and analysis

SMLM image stacks (tiff files) were pre-processed in order to remove background fluorescence using a quantile filter computed on a sliding window. Then, localisations were detected using the algorithm described in [9], based on a wavelet analysis. Subpixel localisation was performed using the radial symmetry center algorithm introduced in [10]. Sample drift was corrected by subtracting the displacement that yielded the best correlation between densities of successive temporal slices grouping 10 000 localisations each. To isolate axons, we applied a Sato filter [11] with a width of 3 pixels on the logarithm of the pixel-wise mean intensity. Then, we used a local thresholding algorithm, provided by [12] to compute a mask over the image. All steps of the analysis were implemented in Python, the code is available at http://gitlab.pasteur.fr. Synapses were manually detoured using an ad-hoc graphic user interface. In total, our analysis includes 321 synapses for which more than 150 trajectories were recorded, coming from 10 different fields of view. A synapse is counted twice if it appears in two or three recordings based on the same field of view but done in different conditions (e.g. some synapses appear in control, KCl and fixed conditions).

In the analysis of experimental trajectories, we considered only trajectories located in the axons, and we split them into two groups: those located outside the synaptic region and those located inside. Synaptic regions were delimited by a density threshold of one tenth of the maximum density of detections in the synapse. The density was estimated using a Gaussian kernel method with a bandwidth of 150 nm. We estimated the apparent effective diffusivity [13] of a trajectory from the sample variance of its single-time-lapse displacements, i.e. 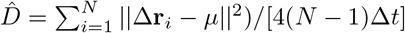, where Δ**r**_*i*_ = **r**_*i*_ − **r**_*i*−1_ is the displacement between the (*i* − 1)th and *i*th recorded positions and 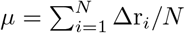 is the average displacement.

### Describing trajectories with latent vectors

The first step of the analysis is to build a latent representation of random walks that does not require strong assumptions about the underlying generative models. In this section, we present the simulation-based inference scheme, the architecture of the neural network used to compute latent vectors from trajectories, and the visualisation of these vectors.

#### Simulation-based inference

In order to ensure that our characterisation of trajectories is accurate, robust and length-independent, it should be trained on an as wide as possible variety of random walks. Hence, we chose to rely on a simulation-based inference procedure [1]. It consists in generating data on which the neural network is subsequently trained. In our case, this amounts to simulating trajectories of a variety of models known to encapsulate different properties of biomolecule dynamics in cells. The physical parameters chosen to simulate these trajectories should at least cover the range of the experimentally observed ones, in order to ensure that the network is able to encode relevant information about the recorded trajectories on which the inference will eventually be performed after its training.

To ensure the diversity of the training set, we simulated trajectories using five different canonical random walk models covering a wide spectrum of possible random walk characteristics:

- the Levy walk (LW) [14–16], which has non-Gaussian increments and exhibits weak ergodicity breaking;
- scaled Brownian motion (sBM) [17–19], which is Gaussian, non-stationary and weakly non-ergodic;
- the Ornstein Uhlenbeck process (OU) [20], a Gaussian, stationary process with exponentially decaying autocorrelations;
- fractional Brownian motion (fBM) [21], which is Gaussian, stationary and exhibits slowly decaying temporal correlations;
- and the continuous time random walk (CTRW) [22, 23], which is non-Gaussian, shows weak ergodicity breaking, ageing, and has discontinuous paths.

The models’ parameters were drawn from the same distributions throughout the entire study and were chosen to cover the entire ranges observed experimentally. Trajectory lengths were drawn from a log-uniform distribution between 7 and 25 points, which corresponds to a mean length of 14 points. The effective diffusivity was drawn from a log-normal distribution with *D*_0_ = 1*μm*^2^*/s*, ⟨log_10_(*D/D*_0_) ⟩ = −0.5 and Var(log_10_(*D/D*_0_)) = 0.5^2^. For the OU model, the relaxation rate *θ* was drawn from …

We added uncorrelated localisation noise to each point of the trajectories, drawn from a centred Gaussian distribution with standard deviation drawn uniformly between 15 and 40 nm (to include the signal intensity dependence of the localisation precision). The time lapse between recordings was set equal to that of the camera (15,4 ms).

The neural network (the architecture of which is detailed below) was then trained to infer two characteristics of interest from the trajectories: their anomalous diffusion exponent (if applicable), and the random walk model from which they were generated among the five described above. Throughout the training, the network processed ∼ 10^6^ independent simulated trajectories.

#### Graph neural network and random walks

Here, we explain how we construct an informative size-independent representation of random walks. We feed trajectories into a neural network, which we train to compute a vector of summary statistics from each trajectory. It contains information relevant to the characterisation of the random walk. Although observed trajectories are not all of the same length, the summary statistics is a vector of constant size. Thus, this vector can be used to compare trajectories of different sizes. Details about the subsequent analyses are found in the next subsections. Here, we focus on the neural network architecture, which is based on our previous work [24]. We refer the interested reader to the Supporting Information for implementation details

Graphical models are methods of choice to handle complex inferences [2, 25], model large scale causal relationships [26] and provide inductive biases in Bayesian inferences [27]. Over the last five years, graph-based analysis methods have been complemented by graph neural networks (GNNs), which extend classical neural network approaches to run directly on graphs. GNNs are efficient at representation learning [28–30] and naturally apply to data of varying dimensions. Furthermore, GNNs are ideally suited to encode natural symmetries of physical systems [31]. For these reasons, we chose to use a such network to process trajectories.

To do so, we represent a trajectory **R** = (**r**_1_, **r**_2_, …, **r**_*N*_) by a directed graph *G* = (*V, E*, **X, Y**). Here *V* = *{*1, 2, …, *N}* are the nodes, each associated with a position in the trajectory, *E* ⊆ *{*(*i, j*)|(*i, j*) ∈ *V* ^2^*}* is the set of edges connecting pairs of nodes, 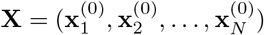 are node feature vectors, and 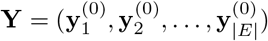 are edge features. Each node feature vector 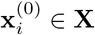, of size *n*_*x*_, captures features associated to the trajectory and to the associated graph. It depends on node *i* and on arbitrary neighborhoods of *i*. The incoming edges of each node connect only to nodes in the past (respecting causality): node *i* receives incoming edges from nodes *i* − Δ_1_, …, *i* − Δ_*max*_, where (Δ_*i*_)_*i*≥1_ is a geometric series. Each edge feature vector 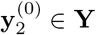, of size *n*_*y*_, contains information about the trajectory’s course between pairs of nodes. The graph construction is illustrated in Fig. 2A.

**Fig 2.**
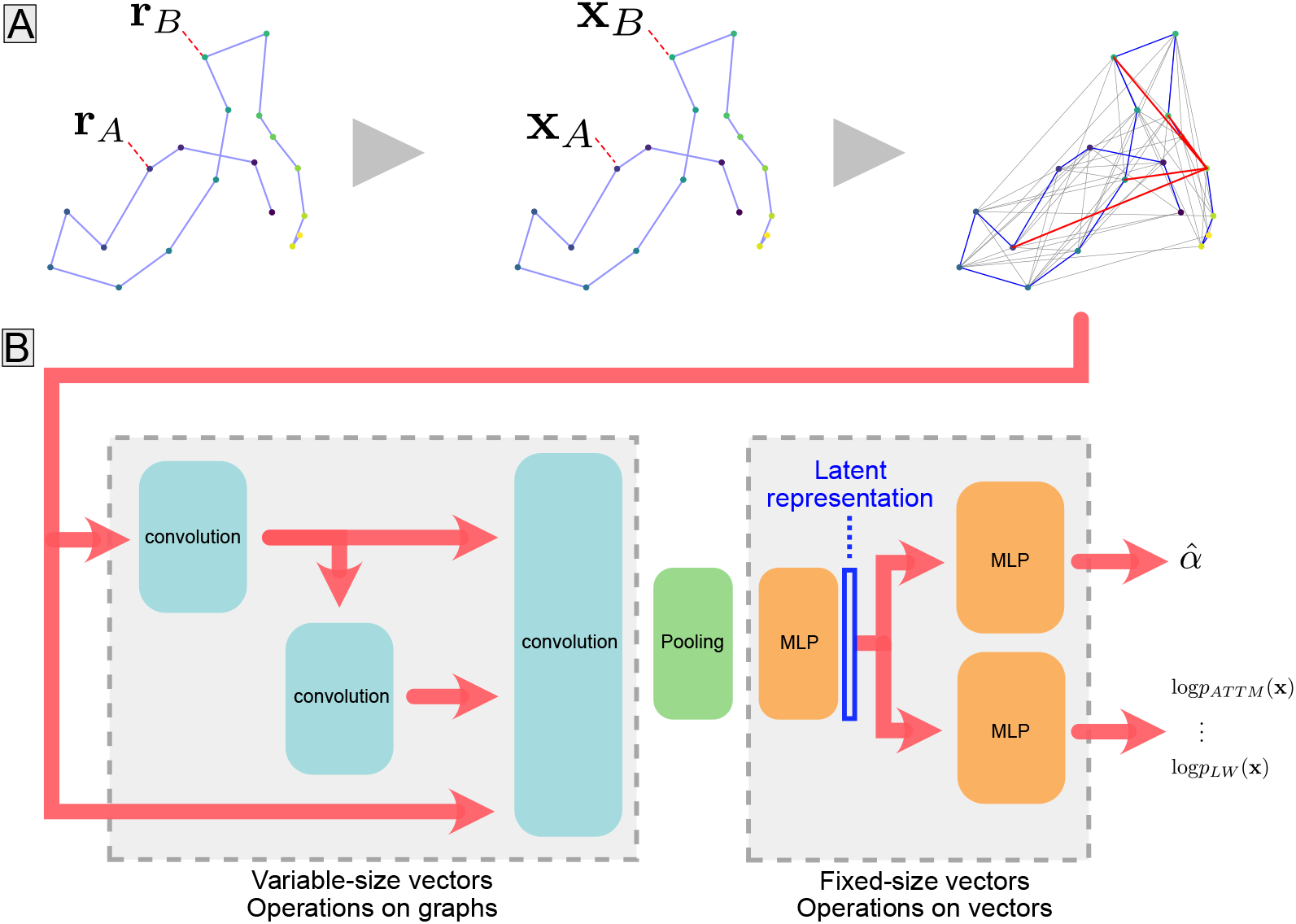
Model architecture. (**A**) Building a trajectory graph. Node and edge features are computed, and edges are drawn between edges following a pre-determined pattern. (**B**) Graph neural network. The graph is passed through a series of graph convolution layers (shown in blue), which propagate information along edges. The pooling operation (green) combines all node feature vectors from a graph into a vector of fixed size representing the graph. This vector is then passed to a multi-layer perceptron (MLP in orange), whose output we refer to as the “latent representation” of the trajectory. The latent representation is fed to two task-specific MLPs: one that predicts the trajectory’s anomalous exponent *α* and one that assigns a vector of probabilities for the trajectory to have been generated by each of the models considered.

A trajectory’s graph and its associated feature vectors are then passed to a series of graph convolution layers, as illustrated in Fig. 2B. Node features vectors are aggregated in the pooling layer such that in the second part of the network, each trajectory is represented by a fixed-size vector. The output of the multi-layer perceptron directly downstream of the pooling layer provides a 16-dimensional vector representing each trajectory, i.e. a summary statistics, which we designate in the following as the “latent representation”, or “latent vector”. During the training phase, the latent vector is fed to two separate MLPs predicting the anomalous exponent and the underlying model of the random walk, respectively. The loss minimised during the network’s training is the sum of two task-specific losses: the mean squared error of the prediction of the anomalous exponent (excluding OU trajectories which are not anomalous random walks) and the cross-entropy of the predicted and true model classes. Training the network on such physically informed tasks makes it build a relevant latent representation of trajectories [24].

#### Latent representation of trajectories

Once processed by the encoder, each trajectory is reduced to a 16-dimensional vector. For visualization purposes, we projected this vector on a 2D plane using a parametric-UMAP to perform the projection [32]. This variation of UMAP allows us to learn the transformation projecting the data from 16 to 2 dimensions solely on simulated trajectories, so that it is independent of the experimental trajectories and is only trained once. The GNN was trained first, then the parametric-UMAP projection was learnt and both their sets of weights were frozen.

We designate as “2D latent representation” the two-dimensional vector, output by the parametric-UMAP, which represents a trajectory in the latent space. Figure 3A shows that latent representations of simulated and experimental data largely overlap, and that the experimental trajectories fall within the region covered by the simulated ones. Figures 3B and 3C show that the random walk model and the diffusivity are prominent determinants of the latent space structure. Figures 3D, 3E and 3F illustrate the diversity of *α*Syn:Eos4 trajectories that can be found in a presynaptic bouton, and how this diversity is captured by the latent representation. Figure 3F highlights the fact that there is a high variability of trajectory dynamics even within a given synapse, suggesting that *α*-synuclein molecules can transition between various dynamic modes.

**Fig 3.**
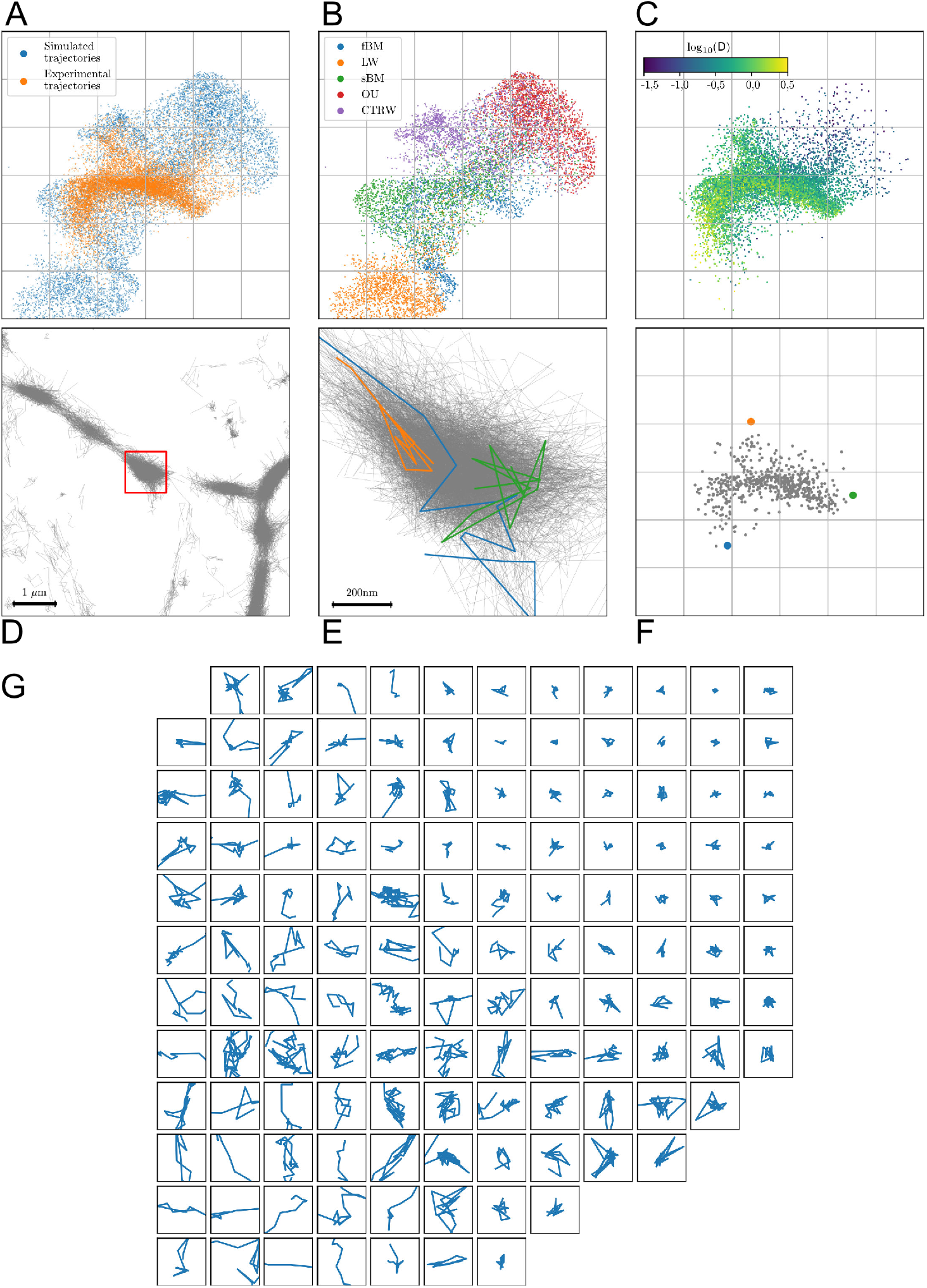
2D Latent space of trajectories. (**A**) Latent representations of simulated versus experimentally recorded trajectories. (**B**) Latent representations of simulated trajectories, coloured by random walk model (fBM: fractional Brownian motion; LW: Levy walk; sBM: scaled Brownian motion; OU: Ornstein-Uhlenbeck process; CTRW: continuous-time random walk). The cropped region at the bottom contains mostly simulated Levy walks. (**C**) Latent representation of experimentally recorded trajectories, coloured according to the estimated log-diffusivity. (**D**) Recorded trajectories at synapses and in the axon. (**E**) Zoom on a presynaptic bouton, delimited by the red square in panel D. Three individual trajectories are highlighted. (**F**) Latent representation of the trajectories at this synapse, with colored dots corresponding to the three trajectories highlighted in E. (**G**) Examples of acquired trajectories, located according to their position in the 2D latent space. Each square has a side length of 1 micrometer.

Using the approach described above, we can associate to any set of trajectories, a set of constant-sized vectors characterising their dynamics. Each microscope recording, or organelle within it, can thus be characterised by a set of *N* 2-dimensional feature vectors, *N* being the number of trajectories. Note that we could have used the latent of vectors of 16 dimensions, but in this application the 2D projection captured enough information about the random walk dynamics. The conditions are thus met to perform statistical testing.

### Statistical testing in the space of latent representations

We develop in this section the statistical test we use to compare the dynamics of single molecules in different organelles and under different biological conditions. Each organelle is characterised by the set of its latent vectors. Thus, we base our statistical test on the comparison of the generating distributions of these vectors. In the absence of *a priori* knowledge of these distributions, we employ a kernel-based approach: the maximum mean discrepancy (MMD) test.

#### Maximum mean discrepancy

Maximum mean discrepancy (MMD), introduced in [4], is a measure of distance between distributions. It was developed to perform statistical testing between two sets of independent observations lying in a metric space *𝒳, X* = *{x*_1_, …, *x*_*m*_*}* drawn from probability measure *p* and *Y* = {*y*_1_, …, *y*_*n*_} drawn from *q*, with the the goal of assessing whether or not *p* and *q* are different.

Given a class ℱ of functions from 𝒳 to ℝ, the MMD between two probability measures *p* and *q* is defined as

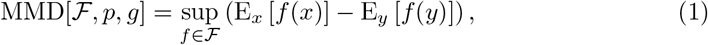

where E_*x*_ and E_*y*_ denote expectation w.r.t. *p* and *q*, respectively.

If the function class is the unit ball in a Reproducing Kernel Hilbert Space (RKHS) [33] ℋ, the square of the MMD can directly be estimated from data samples. Denoting *k* the kernel operator such that ∀*f* ∈ ℱ, *f* (*x*) = ⟨*f, k*(*x*, ·) ⟩, an unbiased estimator of the square of the MMD between *X* and *Y* is given by:

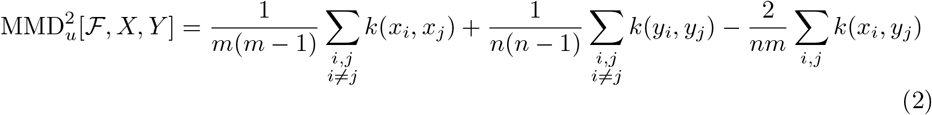

In our case, 𝒳 = ℝ^2^, and we used the classical Gaussian kernel 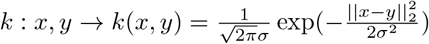. We set the kernel bandwidth *σ* either to the median of the pairwise Euclidian distances between samples from X and Y or we optimised it in specific conditions.

The MMD is capable of detecting subtle differences such as the ones between data generated by generative adversarial networks (GANs) and real data [34]. It has also proved efficient in discovering which variables exhibit the greatest difference between datasets [4, 33].

#### Statistical test

We adapted the bootstrap test described in [4] to assess whether dynamics of *α*-synuclein observed in two experimental conditions exhibit different properties. We denote by *X* and *Y* the two sets of trajectories observed in the two different conditions, drawn from unknown probability densities *p* and *q*. For simplicity, we assume that *X* and *Y* have the same number of elements, *m* = *n*, using the same notation as in the last section. In practice, the number of observed trajectories varies significantly across experimental replicates. To ensure that all replicates have an equal importance when the two sets do not have the same number of trajectories we randomly sub-sampled the larger of the two sets to equalise their sizes. The null hypothesis *H*_0_ of the statistical test is that *p* = *q*, i.e. the two conditions lead to the same distribution of random walks. Under *H*_0_, we approximated the distribution of 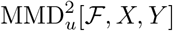 by bootstrapping, i.e. we drew random samples from the union of *X* and *Y* and distributed them in two groups *X′* and *Y ′* (whose sizes respectively match those of *X* and *Y*), on which we computed 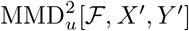. We repeated this procedure a sufficient number of times to obtain an estimation of the distribution of 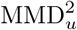 under the assumption that *X′* and *Y′* are drawn from the same distribution. Then, if the original 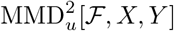 was greater than the 1− *α* quantile of this distribution, we rejected *H*_0_. This test is said to be of level *α*, because with probability *α*, we will reject the null hypothesis when it is actually true.

## Results

### Detecting differences between sets of simulated trajectories

To assess the performance of the full statistical testing framework, we applied it on simulated data. We set the level of statistical significance to *α* = 0.05, and we simulated trajectories as described in Material and Methods.

A first case of our test is to detect changes in the proportions of given types of trajectories between two sets of observations. This is illustrated in Fig. 4, where we show example comparisons in the 2D latent space between two sets with different proportions of their trajectories generated by fBM and sBM. We compared fBM and sBM, since they share numerous features and because for a large range of values of the anomalous exponent they are challenging to distinguish [24]. Furthermore, these two random walk models are highly representative of our experimental data, as can be seen by their latent space occupations (compare Figs. 3A and B).

**Fig 4.**
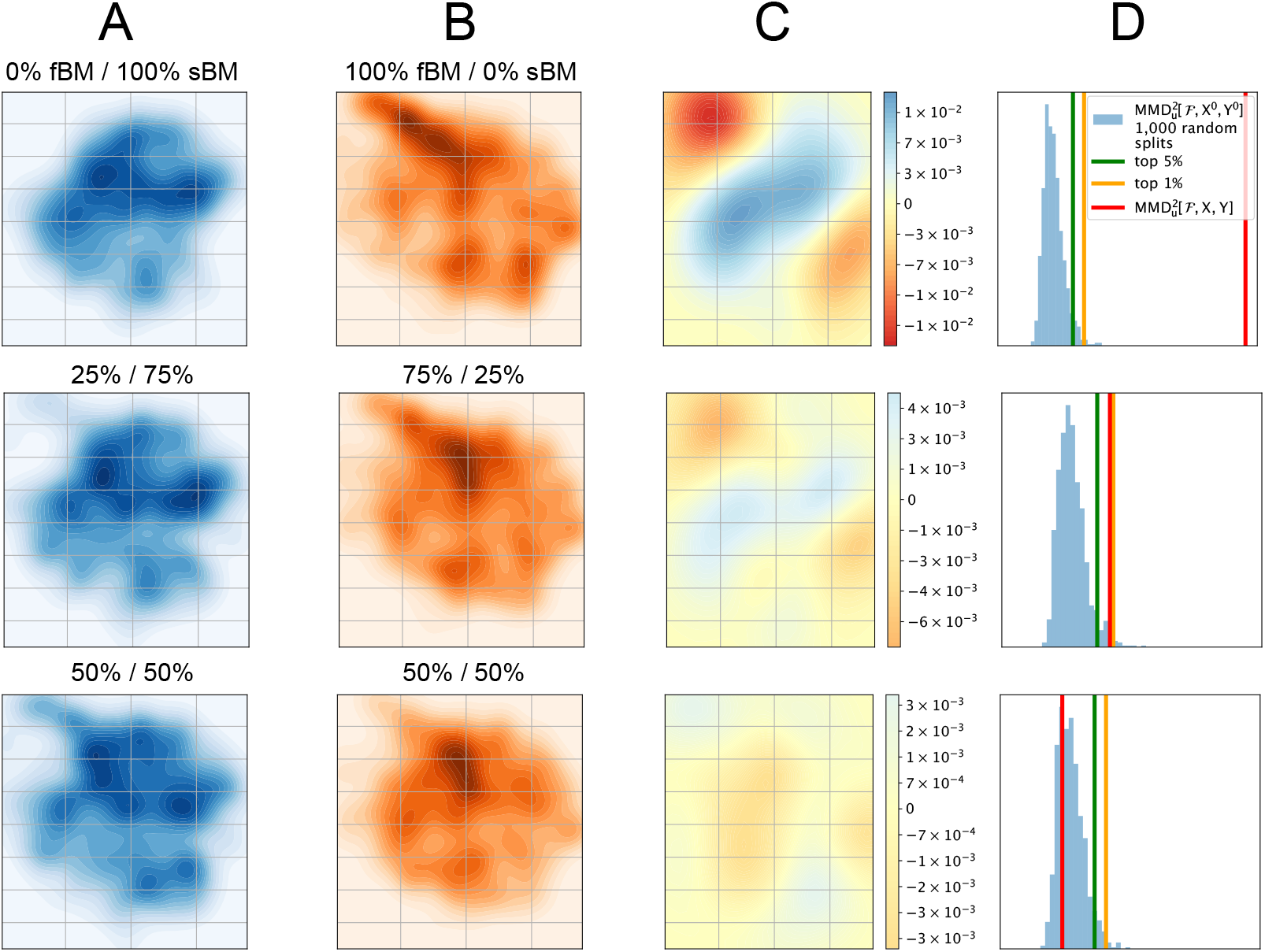
MMD-based statistical test. (**A**), (**B**) Densities of latent vectors in the 2D plane, for two sets of 500 trajectories with different ratios of fBM / sBM trajectories (top: 0% fBM / 100% sBM vs. 100% fBM / 0% sBM, middle: 25% / 75% vs. 75% / 25%, bottom: 50% / 50%). (**C**) Witness functions of the MMD test for difference between A and B, i.e. the function attaining the maximum in Eq. 1, based on the available samples. (**D**) Distribution of the test statistic 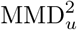 between sets of equal size composed of randomly chosen trajectories of the two sets, with its top 1% (yellow line) and top 5% percentiles (green), as well as the unbiased estimate of the square MMD between the two sets (red).

The difficulty of separating the two populations depends on their relative proportions in the two datasets, and we see that both the amplitude of the witness function (Fig. 4C) as well as the value of the test statistic (Fig. 4D) decrease as the ratio is closer to 1:1. When the two sets are drawn from the same 50/50 distribution, the test does not, and should not, find significant differences between them.

The other main factor determining the difficulty of detecting a difference, is the size of the datasets. Experimentally, changes in biological conditions lead not only to changes in the properties of the random walks. It also leads to changes in the total number of trajectories of a given type. This causes challenges in performing proper statistical testing. To quantitatively assess the effect of both the number of trajectories and the relative proportions belong to different random walk classes, we conducted numerical experiments where we varied these two parameters systematically (Fig. S2A).

Besides differences in the proportions of trajectories generated by different random walk models, the sets may also differ in the models’ parameter values. We thus additionally evaluated the test’s ability to distinguish two sets of fBMs, one with anomalous diffusion exponent *α* = 1 − *δ* and the other with *α* = 1 + *δ*. Our results indicate that in both cases 1 000 trajectories are sufficient to detect subtle changes between distributions (Fig. S2). Fewer trajectories are needed to detect starker differences. In cases where the compared sets are drawn from the same distribution (*ν* = 0 and *δ* = 0), the null hypothesis is rejected in about 5% of cases, consistent with our chosen *α*-level.

One way to further improve these results is to optimise the kernel used to compute the MMD. We show in Fig. S1 how kernel bandwidth and shape affect the power of the test. If the kernel bandwidth is too small, this weakens the test by making it too sensitive to noise. Conversely, if its bandwidth is too large, this prevents the test from detecting subtle changes. We tested the effect of the kernel characteristics in the same setting as illustrated in Figs. 4 and S2A, with *N* = 200 trajectories in each set and comparing sets with 70% fBM / 30% sBM and 30% fBM / 70% sBM. We observed that a Gaussian kernel with radius *σ* equal to the median pairwise distance in the dataset (i.e. *σ* between 1.5 and 2) yields a near-optimal test, in agreement with earlier findings [4]. Finally, while we have here focused on optimizing type II error while controlling type I error (i.e. fixed *α*-level), the parameters and the functional form of the kernel can be adjusted to control either type I or II error while optimising the other [35].

### Differences of *α*-synuclein mobility in axons and at synapses in response to membrane depolarisation

We analysed trajectories of *α*Syn:Eos4 molecules in the axons of cultured cortical neurons and compared them between different subcellular regions, outside or inside the synaptic bouton, and experimental conditions, control, high KCl (leading to synaptic depolarisation) and fixed cells.

The three experimental conditions (control, KCl, fixed) and two subcellular regions (extra- and intra-synaptic) define six populations of trajectories, whose latent space occupation densities are shown in Figs. 5A and 5B. In the remaining four columns of Fig. 5, we illustrate a few comparisons performed between pairs of trajectory populations using our statistical test. Figures 5C and 5D show the latent space densities for the two conditions that are compared in each case, the differences of which are the witness functions shown in Fig. 5E. The distributions of the test statistics under the null hypothesis, obtained after 1 000 bootstrapping iterations, are shown in Fig. 5F, as well as its top 1% and 5% quantiles and the test statistic obtained on the actual two compared populations.

**Fig 5.**
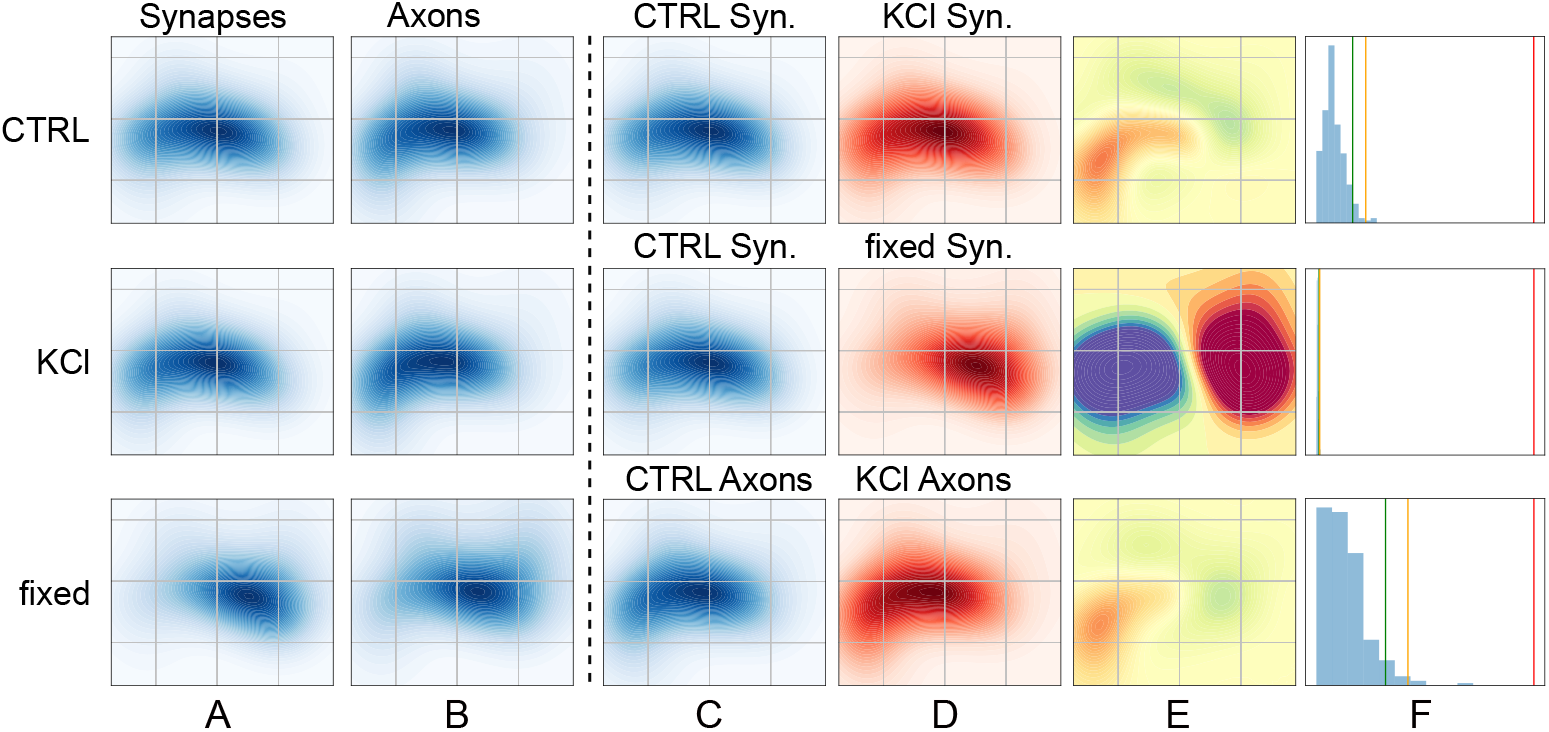
Latent space occupation & statistical testing. Left part: (**A**), (**B**) Latent space occupation densities of *α*Syn:Eos4 trajectories observed in synapses and axons, in the three experimental conditions (control, high KCl, fixed). Right part: each row is one comparison of two sets of trajectories. (**C**), (**D**) Side by side comparison of the latent space occupation densities of the two sets of trajectories used in the comparison in E and F. (**E**) Witness function of the comparison. The colour scale is preserved across rows. (**F**) Histograms illustrating the statistical test based on the MMD. The green and yellow vertical lines represent the top 5% and top 1% quantiles of the null distribution of the squared MMD, respectively, while the red line shows the squared MMD for the experimental data.

In all of the three cases illustrated in Fig. 5, the empirical 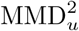 is significantly higher than the top 1% quantile of the null distribution, meaning that our test detects a significant difference in the properties of the trajectories of the two compared subsets at the 1% *α*-level. Note the wide range of magnitudes spanned by these differences, which is well judged by looking at the absolute intensity of the witness functions shown in Fig. 5E. We see that the difference induced by fixation on *α*-synuclein mobility at synapses (Fig. 5E, middle) is much more pronounced than the one induced by high KCl treatment (Fig. 5E, top). This is also apparent when looking at how large 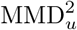 is compared to its distribution under the null hypothesis: in the case of the fixed vs control comparison (Fig. 5F, middle), the histogram of the 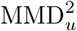 values obtained under the null hypothesis is completely squeezed because the empirical 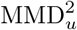 is more than two orders of magnitude larger than the top 1% quantile of the distribution under the null hypothesis. In comparison, KCl treatment produces a less drastic change in *α*-synuclein trajectories located in synaptic terminals (Fig. 5F, top), although this effect is also highly statistically significant (*p* ≪ 0.01).

We further observed that, while their magnitudes differ, the witness functions of the control/KCl comparisons in axons and in synaptic boutons (Fig. 5E, top and bottom) exhibit similar patterns. This indicates that the addition of KCl to the medium can affect the physical properties of many if not all *α*-synuclein molecules in a similar manner, irrespective of their subcellular location. In contrast, *α*-synuclein mobility in fixed neurons appears to be almost entirely abolished, which is seen not only in the amplitude of the change, but also in the fact that the occupation of the latent space displays massive qualitative differences in this condition. This demonstrates that *α*-synuclein is highly mobile in living cells, and helps to put our experimental findings into perspective.

Another feature of the MMD, is the possibility of extracting the points of the feature space that are most important for distinguishing one distribution from another. By finding the local maxima of the *S* statistic, introduced in [36], which is given as the ratio of the mean squared amplitude and the variance of the witness function estimated by bootstrapping, we identify the regions of the latent space where the occupation differs most in the two populations. This enables a straightforward interpretation of the latent space. In Fig. 6, we apply this method to the comparison of intra-synaptic *α*Syn:Eos4 trajectories in the control and KCl conditions. According to this analysis, the representative *α*-synuclein trajectories exhibit a greater mobility in the depolarised state. This is likely the result of a weaker binding of *α*-synuclein at synapses, as reflected in the overall reduction of *α*-synuclein molecules during KCl application (Figs. 1A and 1B).

**Fig 6.**
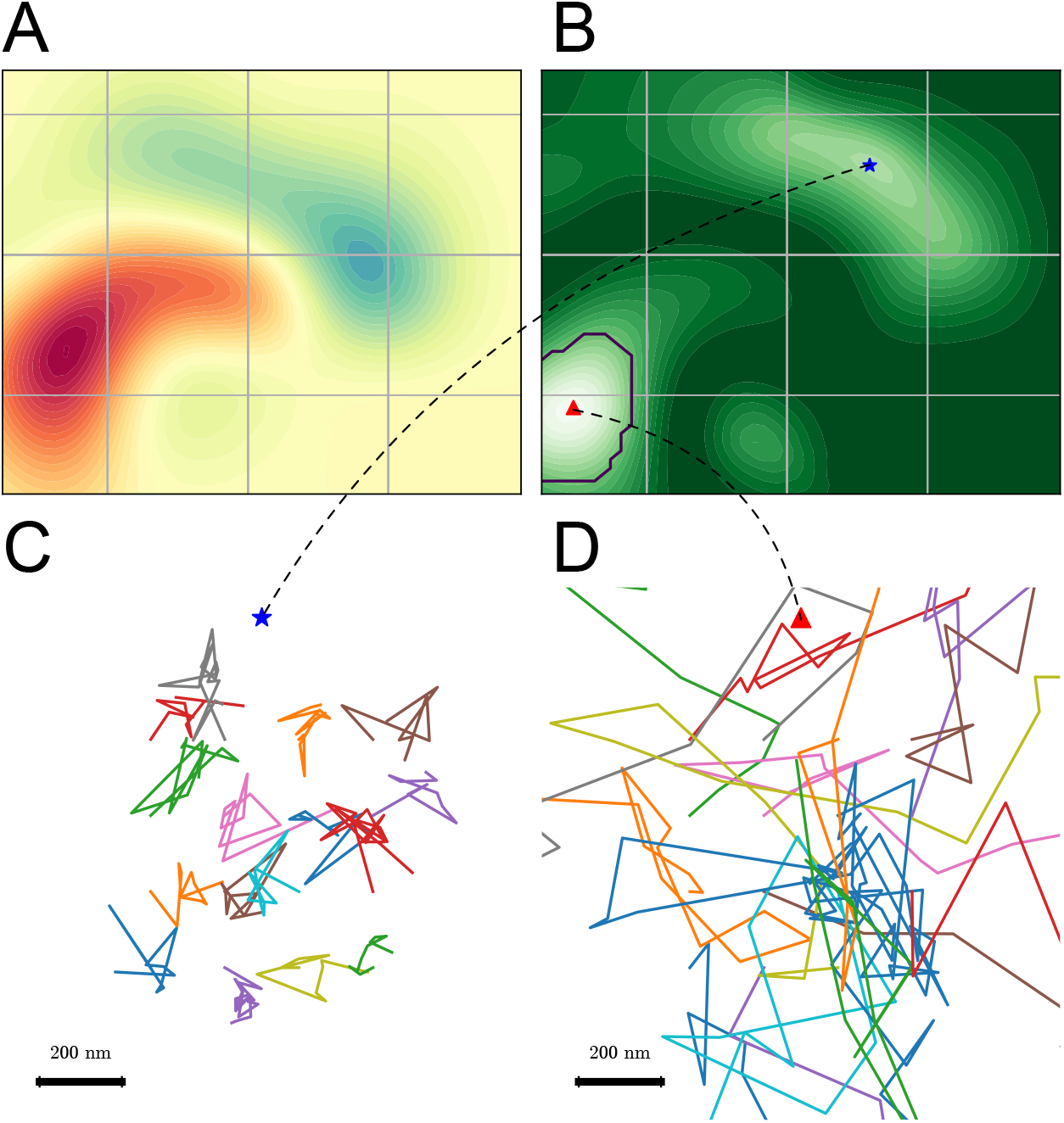
Most salient dynamics. (**A**) Witness function of the comparison of intra-synaptic trajectories, between the control (blue) and KCl (red) conditions. (**B**) Test statistic such as defined in [36], i.e. ratio of the square amplitude divided by the variance. The black contour indicates the “critical region”, *i*.*e*. the region of the latent space most responsible of the statistical difference. (**C**) and (**D**) Illustration, for each maximum of the test statistic, of its 16 closest trajectories.

Furthermore, as illustrated in Fig. S3, we use this statistic to check that all acquired fields of view contribute evenly to the difference between the control and KCl conditions. We looked at the proportion of trajectories coming from each field of view and condition in a region of the latent space which we define as “critical”, based on the value of *S* (it is the contiguous domain containing the maximum of *S* and where *S* is greater than half of its maximum value). We could thus confirm that, on the one hand, trajectories located in this region of the latent space originate from all considered fields of view in a balanced manner, and on the other hand, that within each field of view, the difference of representation of each condition is in the same direction (except for one field of view, where there is almost no difference). This furthermore excludes the possibility that the observed differences are solely due to a single abnormal recording.

### Comparing synapses

The approach we propose is not restricted to inter-condition comparisons, but can be used to compare any two subsets of trajectories. Hence, we can group trajectories by synapse and external condition (control, KCl or fixed), and compute the value of 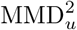 of all the pairs of so-obtained synapses. This provides us with an inter-synapse distance matrix, shown in Fig. 7A. Using these distances, we can embed the trajectory subsets in an Euclidian space, *i*.*e*. summarise each subset by a vector of fixed dimension, using for instance the multi-dimensional scaling (MDS) algorithm [2]. We adapted the MDS algorithm in order to account for the uncertainty that we have in the estimation of the squared distance, which notably depends on the number of observed trajectories per synapse (see Supplementary information). We show in Fig. 7B the vectors obtained when using this method to embed synapses in a 5-dimensional space. The clouds of points corresponding to synapses observed in each condition do not perfectly overlap, which is expected given that we previously showed that conditions were significantly different. However, this view provides an intuitive illustration of the relative extent of inter- and intra-condition variability. More generally, this type of visualisation may be used to illustrate potential continuous shift or clear separations between distinct synaptic regimes observed in different conditions.

**Fig 7.**
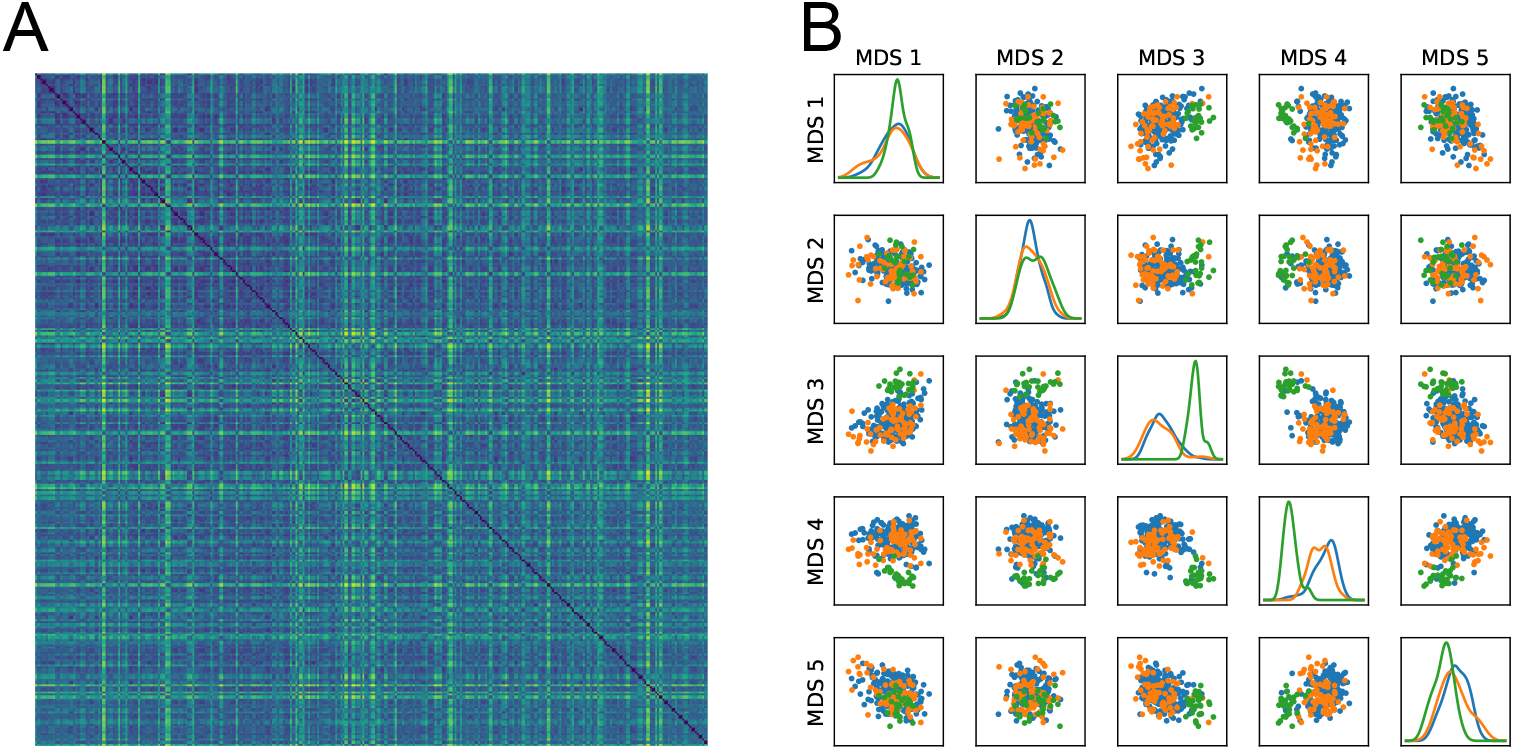
Comparing individual synapses. (**A**) Matrix of the inter-synapse MMD: the value at row *i* and column *j* is colored according to the MMD between trajectories of synapse *i* and synapse *j*. (**B**) 5-dimensional embedding obtained by multi-dimensional scaling (MDS) based on this distance matrix. Dots are coloured according to the condition in which the synapse they represent was observed: CTRL in orange, KCl in blue and fixed in green.

## Discussion

We have introduced a statistical procedure to compare organelles or biological conditions of single molecule experiments. This statistical test does not require explicitly defining the generative models of experimentally recorded random walks. The test consists of two steps. In the first step, an amortised inference is used to reduce any trajectory to a features vector of constant size. In the second, the distribution of features between different conditions is compared by the MMD statistical test.

Our approach allows extracting physically and biologically relevant results without having to assign the biomolecule motion to canonical models. Although these models are instrumental for interpreting the properties of random walks, the complexity and heterogeneity of biological environments at the nanometer/micrometer scale often precludes an unambiguous model assignation.

An alternative approach that would also use a set of canonical random walks as a possible basis for description is Bayesian Model Averaging (BMA). The BMA method does not look for the best model to describe experimental data but rather evaluates the parameters for each of the possible models and then averages the different results. It is challenging to apply this approach to our current problem for two reasons: (i) the space parameters of the different random walks are not identical and (ii) the evaluation of the parameters associated with each random walk by Bayesian methods is not possible for all models. It would be possible to develop Bayesian method variations to estimate the parameters of models that do not have tractable likelihoods. However, the marginalisation of the different models would be computationally intensive. Finally, even though BMA could include information from multiple models, it would still rely on the projection of the experimental date onto a set of canonical models.

We applied our approach to study the dynamics of *α*-synuclein molecules in axons and presynaptic boutons. In agreement with earlier studies of the population dynamics of *α*-synuclein [6, 7], we found that the protein assumes differing dynamic states at synapses and in axons. Depolarisation of the presynaptic terminals through the application of high potassium concentrations shifted the relative frequency of the various states, without necessarily changing the types of diffusion. In other words, our analysis demonstrates clear quantitative changes in the mobility of *α*-synuclein but does not identify qualitative changes due to the extensive overlap in the occupation of the latent space (with the exception of the fixed state).

This statistical testing procedure paves the way to automated analysis of single molecule experiments. Single molecule pharmacology is an emerging field [37, 38], in which the effects of drugs are evaluated at the nanometer scale by studying the spatial properties and dynamics of biomolecules of interest. The possibility to automatically compare different conditions without relying on manually selected generative models of molecule diffusion would be helpful in defining groups of conditions in which a certain effect can be detected. Even though model identification will often be impossible, the properties of the latent space can reveal the source of observed differences. The witness function can thus be instrumental in differentiating changes in the probability of occupancy of specified domains within the latent space between conditions. As illustrated on Fig. 6, going from a region of the latent space to an intelligible trait of trajectories is rather intuitive, hence the interest of this method to orient further analysis and build biologically relevant hypotheses.

Beyond the automation of the analysis procedure between biological conditions, our approach is well suited for exploratory data analysis. The capacity to project individual, differently sized trajectories into finite sized vectors makes it possible to study precise sub-cellular compartments or organelles in a standardised form, and thus allows to test statistical differences between these regions. Hence, recorded single molecule data can be searched in order to detect and characterise regions of the cell that have different statistical properties. This exploration can be done even in regions with different trajectory densities, as is the case for *α*-synuclein at synapses versus axonal domains.

One of the current limitations of the current approach is the difficulty in evaluating the type II error [35] bounds on the statistical test. The MMD test is applied within the latent space of the GNN. This manifold is built using a set of non-linear operations, which depends both on the numerical trajectories seen during the training and on the cost function being optimised. Hence, there may be domains within the latent space that could lead to improper sensitivity of the statistical test. As can be seen in [24] and in Fig. 3, different types of random walks occupy domains of different size and there is a large overlap of the regions. Since our approach relies on a simulation based framework, it is possible to use numerical simulations matching the experimental occupancy of the latent space to evaluate the accuracy of the test. Furthermore, extensive simulations and check if the statistical test misbehaves, even though this procedure can be time consuming. In order to further improve the statistical power of the test, one could optimise the kernel with which the MMD is computed. Along these lines, a possible variant of this method could rely on an encoder network trained not on a supervised inference task but rather to maximise the MMD between two sets of experimentally recorded trajectories. This, however, would require substantially a larger quantity of experimental data.

## Acknowledgments

We thank Alexandre Blanc, Michael Lelek, Sylvain Prigent, Mohamed El Beheiry, Srini Turaga, Raphael Voituriez & Bassam Hajj for helpful discussions. Fumihiro Niwa is thanked for technical help. This study was funded by the Institut Pasteur, *L’Agence Nationale de la Recherche* (TRamWAy, ANR-17-CE23-0016 to JBM), the INCEPTION project (PIA/ANR-16-CONV-0005, OG), and the *“Investissements d’avenir”* programme under the management of Agence Nationale de la Recherche, reference ANR-19-P3IA-0001 (PRAIRIE 3IA Institute) to JBM & CLV.

The funding sources had no role in study design, data collection and analysis, decision to publish, or preparation of the manuscript.

## Supporting information

### Multi-dimensional scaling with uncertainty on estimated distances

When estimating 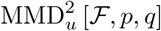 from sets of trajectories *X* ∼ *p* and *Y* ∼ *q*, the uncertainty directly depends on the number of samples in both *X* and *Y*. In our case, the uncertainty, which can be evaluated by bootstrapping, is sometimes of the same order of magnitude than the estimated value. Furthermore, the number of elements per set (here, the number of trajectories per synapse), spans more than an order of magnitude and uncertainty thus greatly varies from one measure to the other. This should be taken into account when using 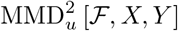 to embed subsets of trajectories.

Hence, starting from a matrix of squared distances **D**^2^ between *N* sets of trajectories, and a matrix of uncertainties of these squared distances 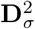, we obtain a set of *N* Euclidian vectors {**x**_1_, …, **x**_*N*_} by maximizing the probability of the resulting squared distances, assuming that they follow Gaussian laws whose means are the coefficients of **D**^2^ and standard deviations coefficients of 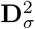. This amounts to solving the following optimisation problem:

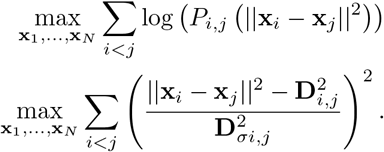

We do so using a gradient ascent method.

### Graph neural network features and architecture

#### Node and edge features

Prior to entering the encoder, each trajectory is turned into a graph as described in. To each node is associated a vector containing the following features:

- the normalised time: *i/N*;
- the cumulative distance covered by the trajectory up to *i*: ∑_*k*≤*i*_‖Δ**r**_*k*_‖_2_;
- the cumulative squared distance covered by the trajectory up to 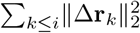;
- the maximum step size up to *i*: max_*k*≤*i*_‖Δ**r**_*k*_‖_2_.

Similarly, each edge is associated to the following set of features:

- the normalised time difference: (*j* − *i*)*/N*;
- the distance: ‖**r**_*j*_ − **r**_*i*_‖_2_;
- the dot product of jumps: 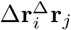 (equal to Δ**r**_*i*_Δ**r**_*j*_ for 1D trajectories);
- the distance covered by the trajectory between *i* and *j*: ∑_*i<k*≤*j*_ ‖Δ**r**_*k*_ ‖_2_ =∑ _*k*≤*j*_ ‖Δ**r**_*k*_ ‖_2_ − ∑_*k*≤*i*_‖Δ**r**_*k*_ ‖_2_
- sum of square step sizes between *i* and *j*: 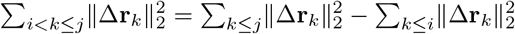.

Features based on distances are computed on a normalised three different versions of the trajectories, corresponding to three different normalisation factors: the total covered distance, the standard deviation of the positions, and the standard deviation of step sizes. These three scales are concatenated to the output of the pooling layer and are thus processed by the perceptron which produces the latent representation of a trajectory (see Fig. 2).

#### GNN architecture

The architecture of the GNN used in the summary network is similar to the encoder network proposed in [24], with the difference that we here additionally apply edge features. Node and edge features are first passed to perceptrons, which embeds them in a homogeneous space. The network is then composed of three successive convolution layers (one taken from [39] and two edge-conditioned layers taken from [40]) outputting node features matrices **x**^(1)^, **x**^(2)^ and **x**^(3)^, each of 32 dimensions, which are summed to form **x**^(*f*)^. The rows of this matrix of nodes features are then averaged during the pooling step, to keep just one row per graph, i.e., per trajectory. This vector is subsequently passed to a three-layer perceptron, the output of which is the summary statistics vector. All multi-layer perceptrons have a leaky-ReLU activation with slope 0.1 for negative values. We summarise their shapes in table 1

**Table 1.** Shapes of multi-layer perceptrons used in the network.

| MLP | Layers |
| --- | --- |
| edge features embedding | (13,32,32,16) |
| node features embedding | (10,32,32,10) |
| edge convolution (x2) | (16,32,32,1024) |
| final embedding | (38,32,16,16) |
| alpha predictor | (16,128,64,64,16,1) |
| model classifier | (16,32,16,5) |

**Fig S1.**
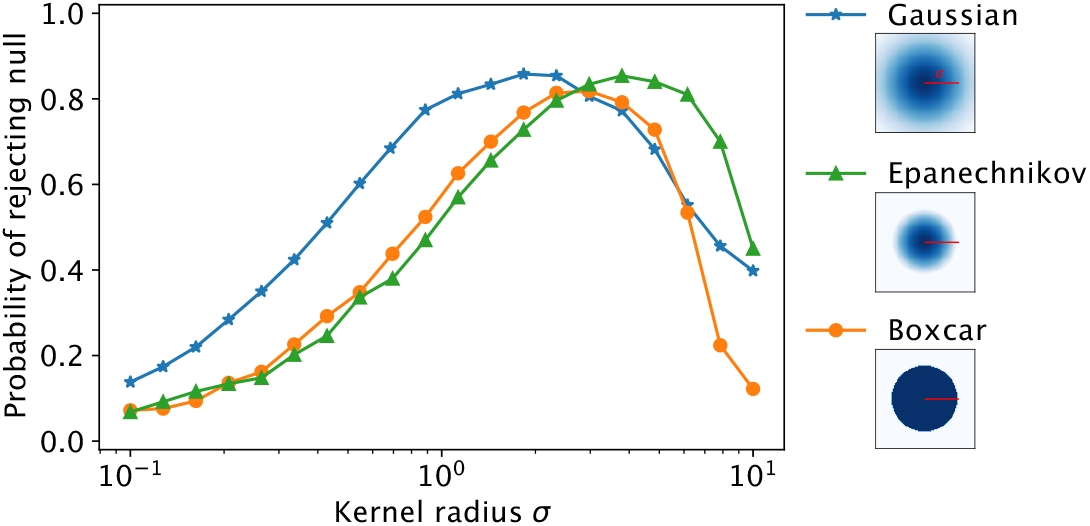
Influence of kernel parameters on the test’s power. Probability of rejecting the null hypothesis that the two sets of trajectories are drawn from the same distribution, with varying kernel types and radii. The sets are drawn with the same characteristics as those used for Fig. S2A with *N* = 200 and *ν* = 0.2.

**Fig S2.**
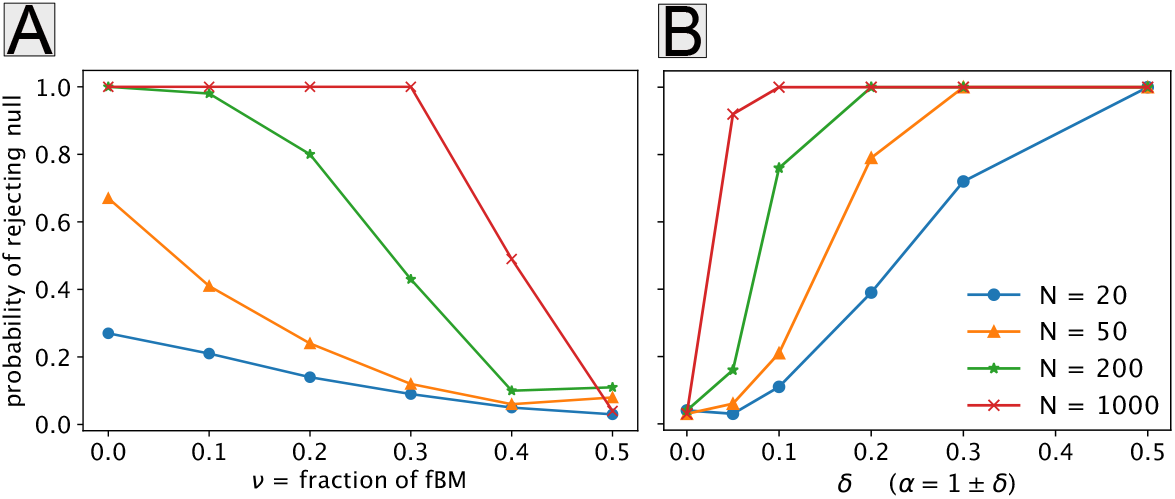
Performance of the statistical test on simulated trajectories. **A**: Probability of detecting a difference between two sets of *N* trajectories composed of a fraction *ν* of fractional Brownian motions and 1 − *ν* of scaled Brownian motions. **B**: Probability of detecting a difference between two sets of *N* fractional Brownian motions, one with anomalous diffusion exponent *α* = 1 − *δ* and the other with *α* = 1 + *δ*. Probabilities are estimated by performing the test 100 times; at each trial, new trajectories are simulated and bootstrap-estimation of the distribution of 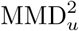 under the null hypothesis is done using 100 random splits.

**Fig S3.**
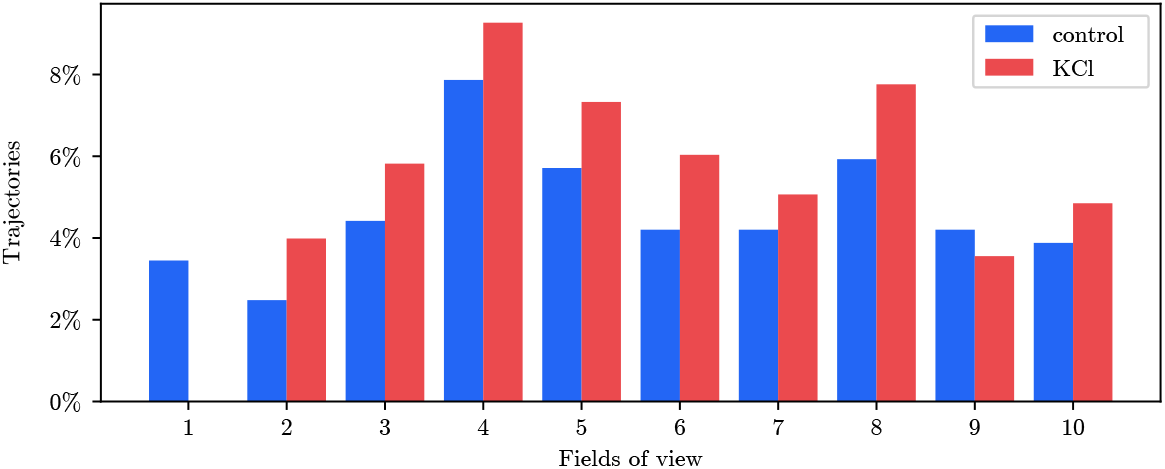
Origin of trajectories found in the critical region. These counts were obtained using a set composed of the same number *n* = 1, 000 of trajectories from each microscopy recording. We randomly subsampled those who had more intra-synaptic trajectories, and discarded those who had less than 1,000 (hence the column with missing number of KCl trajectories).

## References

1. Cranmer K, Brehmer J, Louppe G. The frontier of simulation-based inference;117(48):30055–30062. doi:10.1073/pnas.1912789117.

2. Bishop CM. Pattern Recognition and Machine Learning. Softcover reprint of the original 1st ed. 2006 edition ed. Springer;.

3. LeCun Y, Bengio Y, Hinton G. Deep learning;521(7553):436–444. doi:10.1038/nature14539.

4. Gretton A, Borgwardt KM, Rasch MJ, Schölkopf B, Smola A. A Kernel Two-Sample Test;13(25):723–773.

5. Specht CG. A Quantitative Perspective of Alpha-Synuclein Dynamics–Why Numbers Matter. Frontiers in Synaptic Neuroscience. 2021;13.

6. Spinelli KJ, Taylor JK, Osterberg VR, Churchill MJ, Pollock E, Moore C, et al. Presynaptic alpha-synuclein aggregation in a mouse model of Parkinson’s disease. Journal of Neuroscience. 2014;34(6):2037–2050.

7. Fortin DL, Nemani VM, Voglmaier SM, Anthony MD, Ryan TA, Edwards RH. Neural activity controls the synaptic accumulation of α-synuclein. Journal of Neuroscience. 2005;25(47):10913–10921.

8. Ludwig A, Serna P, Morgenstein L, Yang G, Bar-Elli O, Ortiz G, et al. Feasibility analysis of semiconductor voltage nanosensors for neuronal membrane potential sensing;. Available from: https://www.biorxiv.org/content/10.1101/838342v1.

9. Izeddin I, Boulanger J, Racine V, Specht C, Kechkar A, Nair D, et al. Wavelet analysis for single molecule localization microscopy. Optics express. 2012;20(3):2081–2095.

10. Parthasarathy R. Rapid, accurate particle tracking by calculation of radial symmetry centers. Nature methods. 2012;9(7):724–726.

11. Sato Y, Nakajima S, Shiraga N, Atsumi H, Yoshida S, Koller T, et al. Three-dimensional multi-scale line filter for segmentation and visualization of curvilinear structures in medical images. Medical image analysis. 1998;2(2):143–168.

12. Van der Walt S, Schönberger JL, Nunez-Iglesias J, Boulogne F, Warner JD, Yager N, et al. scikit-image: image processing in Python. PeerJ. 2014;2:e453.

13. Vestergaard CL, Blainey PC, Flyvbjerg H. Optimal Estimation of Diffusion Coefficients from Single-Particle Trajectories;89(2):022726. doi:10.1103/PhysRevE.89.022726.

14. Klafter J, Zumofen G. Lévy statistics in a Hamiltonian system. Physical Review E. 1994;49(6):4873.

15. Palyulin VV, Chechkin AV, Metzler R. Lévy flights do not always optimize random blind search for sparse targets. Proceedings of the National Academy of Sciences. 2014;111(8):2931–2936.

16. Koren T, Lomholt MA, Chechkin AV, Klafter J, Metzler R. Leapover lengths and first passage time statistics for Lévy flights. Physical review letters. 2007;99(16):160602.

17. Lim S, Muniandy S. Self-similar Gaussian processes for modeling anomalous diffusion. Physical Review E. 2002;66(2):021114.

18. Jeon JH, Chechkin AV, Metzler R. Scaled Brownian motion: a paradoxical process with a time dependent diffusivity for the description of anomalous diffusion. Physical Chemistry Chemical Physics. 2014;16(30):15811–15817.

19. Sposini V, Metzler R, Oshanin G. Single-trajectory spectral analysis of scaled Brownian motion. New Journal of Physics. 2019;21(7):073043.

20. Crispin G. Handbook of Stochastic Methods: for Physics, Chemistry and natural sciences. 4th ed. Springer;.

21. Mandelbrot BB, Van Ness JW. Fractional Brownian motions, fractional noises and applications. SIAM review. 1968;10(4):422–437.

22. Scher H, Montroll EW. Anomalous transit-time dispersion in amorphous solids. Physical Review B. 1975;12(6):2455.

23. Magdziarz M, Weron A, Burnecki K, Klafter J. Fractional Brownian motion versus the continuous-time random walk: A simple test for subdiffusive dynamics. Physical review letters. 2009;103(18):180602.

24. Verdier H, Duval M, Laurent F, Cassé A, Vestergaard CL, Masson JB. Learning physical properties of anomalous random walks using graph neural networks. Journal of Physics A: Mathematical and Theoretical. 2021;54(23):234001.

25. Yedidia JS. Message-Passing Algorithms for Inference and Optimization;145(4):860–890. doi:10.1007/s10955-011-0384-7.

26. Koller D. Probabilistic Graphical Models: Principles and Techniques (Adaptive Computation and Machine Learning series). The MIT Press;.

27. Lake BM, Salakhutdinov R, Tenenbaum JB. Human-level concept learning through probabilistic program induction. Science. 2015;350(6266):1332–1338.

28. Fey M, Lenssen JE. Fast Graph Representation Learning with PyTorch Geometric. In: ICLR Workshop on Representation Learning on Graphs and Manifolds;.

29. Kipf TN, Welling M. Semi-Supervised Classification with Graph Convolutional Networks;.

30. Qi CR, Su H, Mo K, Guibas LJ. PointNet: Deep Learning on Point Sets for 3D Classification and Segmentation;.

31. Satorras VG, Hoogeboom E, Welling M. E(n) Equivariant Graph Neural Networks;.

32. Sainburg T, McInnes L, Gentner TQ. Parametric UMAP embeddings for representation and semi-supervised learning;.

33. Gretton A. Reproducing Kernel Hilbert Spaces in Machine Learning; p. 133.

34. Sutherland DJ, Tung HY, Strathmann H, De S, Ramdas A, Smola A, et al. Generative Models and Model Criticism via Optimized Maximum Mean Discrepancy;.

35. Wasserman L. Hypothesis Testing and p-values. In: Wasserman L, editor. All of Statistics: A Concise Course in Statistical Inference. Springer Texts in Statistics. Springer;. p. 149–173. Available from:https://doi.org/10.1007/978-0-387-21736-9_10.

36. Jitkrittum W, Szabó Z, Chwialkowski KP, Gretton A. Interpretable Distribution Features with Maximum Testing Power. In: Advances in Neural Information Processing Systems. vol. 29. Curran Associates, Inc.;.Available from: https://proceedings.neurips.cc/paper/2016/hash/0a09c8844ba8f0936c20bd791130d6b6-Abstract.html.

37. Ruan Q, Macdonald PJ, Swift KM, Tetin SY. Direct single-molecule imaging for diagnostic and blood screening assays;118(14):e2025033118. doi:10.1073/pnas.2025033118.

38. Gunnarsson A, Snijder A, Hicks J, Gunnarsson J, Höök F, Geschwindner S. Drug Discovery at the Single Molecule Level: Inhibition-in-Solution Assay of Membrane-Reconstituted -Secretase Using Single-Molecule Imaging;87(8):4100–4103. doi:10.1021/acs.analchem.5b00740.

39. Kipf TN, Welling M. Semi-Supervised Classification with Graph Convolutional Networks;.

40. Simonovsky M, Komodakis N. Dynamic edge-conditioned filters in convolutional neural networks on graphs. In: Proceedings of the IEEE conference on computer vision and pattern recognition; 2017. p. 3693–3702.

